# Co-infection of influenza A virus enhances SARS-CoV-2 infectivity

**DOI:** 10.1101/2020.10.14.335893

**Authors:** Lei Bai, Yongliang Zhao, Jiazhen Dong, Simeng Liang, Ming Guo, Xinjin Liu, Xin Wang, Zhixiang Huang, Xiaoyi Sun, Zhen Zhang, Lianghui Dong, Qianyun Liu, Yucheng Zheng, Danping Niu, Min Xiang, Kun Song, Jiajie Ye, Wenchao Zheng, Zhidong Tang, Mingliang Tang, Yu Zhou, Chao Shen, Ming Dai, Li Zhou, Yu Chen, Huan Yan, Ke Lan, Ke Xu

## Abstract

The upcoming flu season in the northern hemisphere merging with the current COVID-19 pandemic raises a potentially severe threat to public health. Through experimental co-infection of IAV with either pseudotyped or SARS-CoV-2 live virus, we found that IAV pre-infection significantly promoted the infectivity of SARS-CoV-2 in a broad range of cell types. Remarkably, increased SARS-CoV-2 viral load and more severe lung damage were observed in mice co-infected with IAV *in vivo*. Moreover, such enhancement of SARS-CoV-2 infectivity was not seen with several other viruses probably due to a unique IAV segment as an inducer to elevate ACE2 expression. This study illustrates that IAV has a special nature to aggravate SARS-CoV-2 infection, and prevention of IAV is of great significance during the COVID-19 pandemic.

## Introduction

The outbreak of Severe Acute Respiratory Syndrome Coronavirus 2 (SARS-CoV-2) at the end of 2019 has become pandemic worldwide. Up to date, there had been more than 36 million confirmed infected cases and 1 million deaths globally (https://covid19.who.int/). The ending time and the final severity of the current COVID-19 pandemic wave are still uncertain. Meanwhile, the upcoming seasonal influenza merging with the current pandemic might bring more challenges and pose a bigger threat to public health. There are many debates on whether seasonal flu would impact the severity of the COVID-19 pandemic and whether massive influenza vaccination is necessary for the coming winter. However, no experimental evidence is available concerning IAV and SARS-CoV-2 co-infection.

It is well known that disease symptoms from SARS-CoV-2 and IAV infections are quite similar, such as fever, cough, pneumonia, acute respiratory distress syndrome, etc(*1, 2*). Moreover, both SARS-CoV-2 and IAV are airborne transmitted pathogens that infect the same human tissues such as the respiratory tract, nasal, bronchial, and alveolar epithelial cultures(*3, 4*). Besides, alveolar type II cells (AT2 pneumocytes) appeared to be preferentially infected by SARS-CoV-2, which were also the primary site of IAV replication(*5, 6*). Therefore, the overlap of the COVID-19 pandemic and seasonal influenza would pose a large population under the high risks of co-occurrent infection by these two viruses(*7*).

Unfortunately, during the last winter flu season in the southern hemisphere, there was little epidemiological evidence about the interaction between COVID-19 and flu, probably due to a low IAV infection rate resulted from social distancing(*8, 9*). A case report showed that three out of four SARS-CoV-2 and IAV co-infected patients rapidly develop to respiratory deterioration(*10*). On the contrary, other reports only observed mild symptoms in limited co-infection outpatients(*11*). Thus, the clinical co-infection outcomes are still unclear when a large population will face the threats of both viruses.

In this study, we tested whether IAV infection could affect the subsequent SARS-CoV-2 infection in both infected cells and mice. The results demonstrate that the pre-infection of IAV strongly enhances the infectivity of SARS-CoV-2 by boosting viral entry in the cells and by elevating viral load plus more severe lung damage in infected mice. These data suggest a clear auxo-action of IAV on SARS-CoV-2 infection, which implies the great importance of influenza virus and SARS-CoV-2 co-infection to public health.

## Results

### IAV promotes SARS-CoV-2 virus infectivity

To study the interaction between IAV and SARS-CoV-2, A549 (a hypotriploid alveolar basal epithelial cell line) cells that are susceptible to IAV infection but usually do not support SARS-CoV-2 infection were applied to test whether IAV pre-infection would modulate the infectivity of SARS-CoV-2. Pseudotyped VSV luciferase-reporter particles bearing SARS-CoV-2 spike protein (pSARS-CoV-2) were used to reflect the virus entry activity(*12*). The cells were firstly infected with IAV (A/WSN/1933[H1N1]) or mock-infected for 6 h, 12 h, or 24 h respectively, and then infected with the pSARS-CoV-2 virus for another 24 h (experimental scheme shown in Fig.1A). The data in Fig. 1B showed that A549 was converted to be highly sensitive (up to 10,000-fold) against the pSARS-CoV-2 virus after different doses of IAV infections (from low MOI of 0.01 to high MOI of 1, also shown by pSARS-CoV-2 with mCherry reporter in Fig. S1). In contrast, the pre-infection of IAV had no impacts on pseudotyped VSV particles bearing VSV-G protein (Fig.1C). We further tested more cell lines to show that the enhancement of the pSARS-CoV-2 infectivity by IAV was a general effect although the increased folds were different (lower basal level of infectivity, higher enhancement fold) (Fig.1D).

**Fig. 1.**
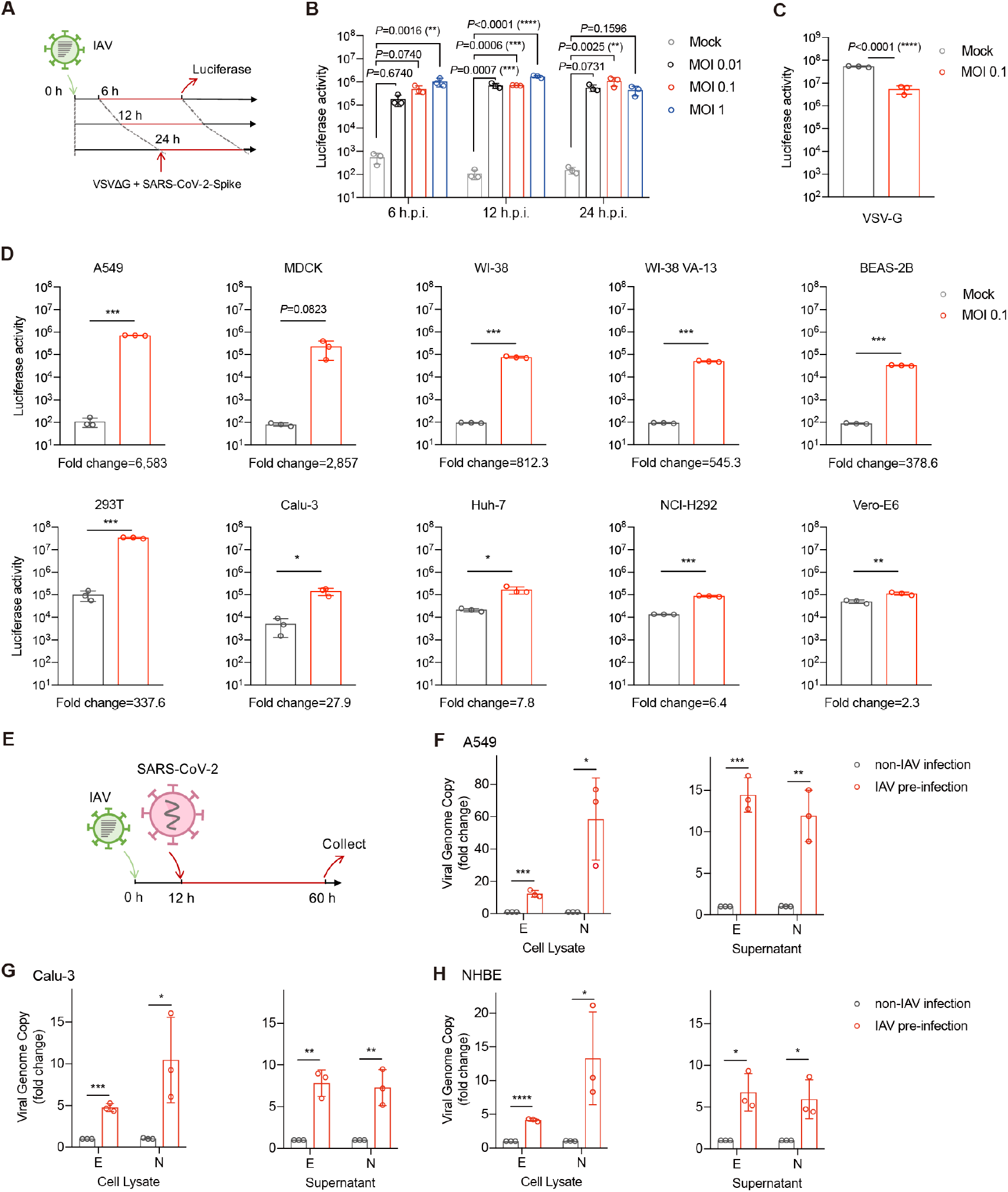
IAV promotes SARS-CoV-2 virus infectivity. (**A**) Diagram of the experimental procedure. (**B**) A549 cells were infected with A/WSN/33 at indicated MOIs. At 6, 12, 24 hours post-IAV-infection, cells were infected with pSARS-CoV-2 for another 24 hours. Luciferase activity was measured to reflect virus entry efficiency. P values are from unpaired One-way ANOVA. (**C**) A549 cells were infected with A/WSN/33 at MOI 0.1. At 12 hours post-IAV-infection, cells were infected with VSV-G-Luc for another 24 hours. Luciferase activity was measured to reflect virus entry efficiency. (**D**) The indicated cells were infected with WSN at MOI 0.1. At 12 hours post-IAV-infection, cells were infected with pSARS-CoV-2 for another 24 hours. Luciferase activity was measured to reflect virus entry efficiency. (**E**) The experimental procedure of IAV and live SARS-CoV-2 co-infection. A549 (**F**), Calu-3 (**G**), and NHBE (**H**) cells were pre-infected with WSN at MOI 0.1 for 12 hours. Cells were then infected with live SARS-CoV-2 at MOI 0.01 for another 48 hours. Total RNA in cell lysates and the supernatant were collected to detect E and N gene by Taqman-qRT-PCR. The data were expressed as fold changes of viral RNA levels in IAV pre-infection cells relative to the non-IAV infection control. Values are mean ± s.d. of three independent results. *P≤0.05, **P≤0.01, ***P≤0.001, ****P≤0.0001.

To validate the above results, we substituted the pSARS-CoV-2 with the SARS-CoV-2 live (experimental scheme shown in Fig.1E). We found that the pre-infection of IAV strongly increased the copy numbers of the SARS-CoV-2 genome (E and N genes) in both cell lysates and supernatants of A549 (~15 folds) (Fig.1F). Notably, in Calu-3 (Fig.1G) and NHBE (Fig.1H) cells that are initially susceptible to SARS-CoV-2, IAV pre-infection could further increase >5 folds of SARS-CoV-2 infectivity.

Collectively, these data suggest an auxo-action of IAV on SARS-CoV-2 in a broad range of cell types.

### IAV and SARS-CoV-2 co-infection in mice results in increased SARS-CoV-2 viral load and more severe lung damage

The hACE2 transgenic mice were applied to study the interaction between IAV and SARS-CoV-2 *in vivo*. Mice were infected with 3×10^5^ PFU of SARS-CoV-2 with or without 2000 PFU of IAV pre-infection and were then sacrificed two days later after SARS-CoV-2 infection (the experimental scheme is shown in Fig. 2A). The viral RNA genome copies from lung homogenates confirmed that SARS-CoV-2 efficiently infected both groups (more than 4×10^8^ N gene copies) (Fig. 2B), while the influenza NP gene was only detected in IAV pre-infection group (Fig. 2B). Intriguingly, a significant increase in SARS-CoV-2 viral load (12.9-fold increase in E gene and 6.6-fold increase in N gene) was observed in lung homogenates from co-infection mice compared to that from SARS-CoV-2 single-infected mice (Fig. 2C). The histological data in Fig. 2D further illustrated that IAV and SARS-CoV-2 co-infection induced more severe lung pathologic changes with massive infiltrating cells and obvious alveolar necrosis as compared to SARS-CoV-2 single infection or mock infection.

**Fig. 2.**
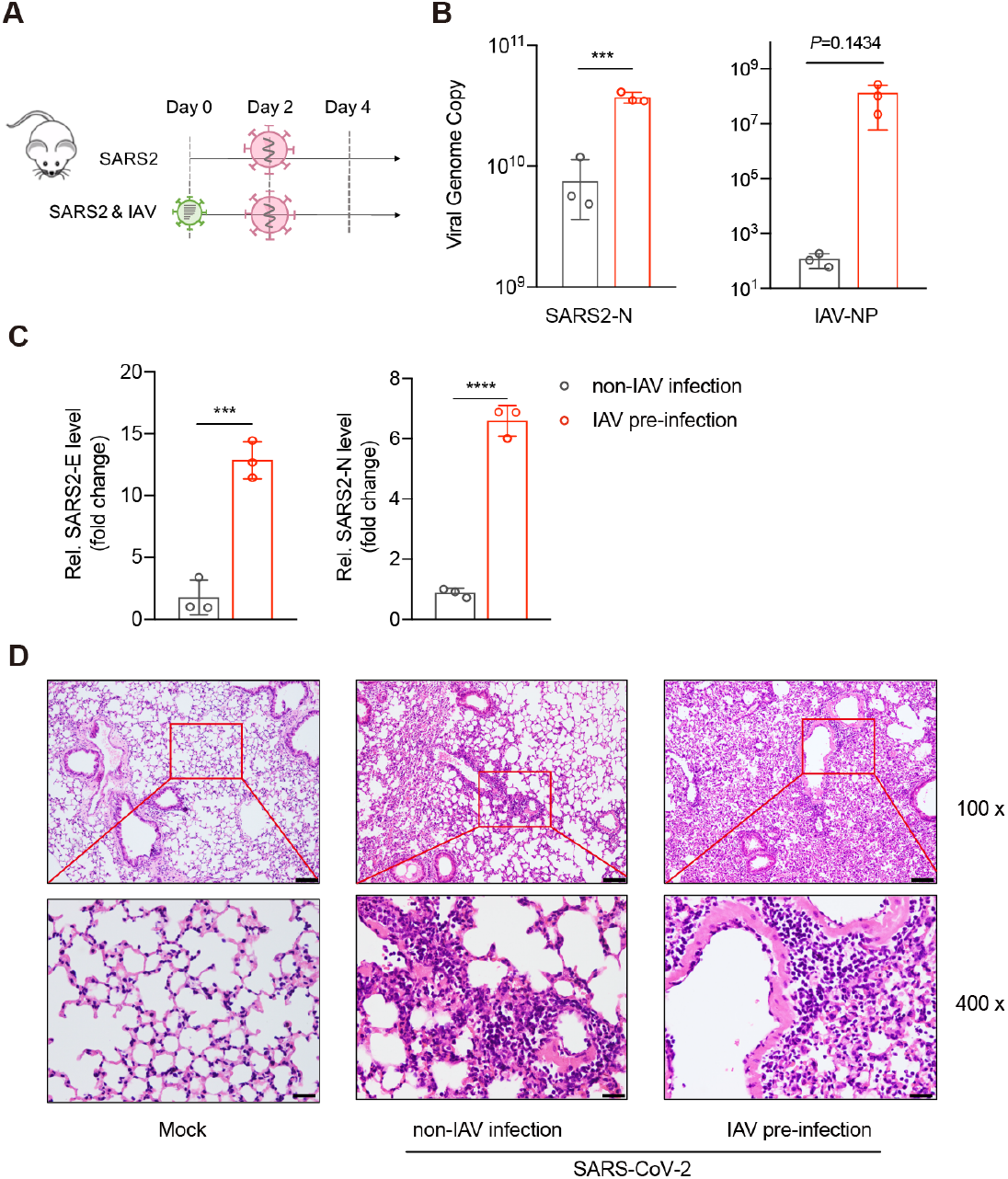
IAV and SARS-CoV-2 co-infection induce more severe pathology in infected mice. (**A**) Diagram of the experimental procedure. K18 hACE2 transgene mice were firstly intranasally infected with 2000 PFU of WSN or PBS at Day 0. Two days post-IAV-infection, mice were intranasally infected with 3 x10^5^ live SARS-CoV-2 or PBS. At day 4, half lung tissues of all the mice were homogenized to detect RNA or protein levels. (**B**) The quantitative viral genome copy numbers of SARS-CoV-2 N (**B, left**) or IAV NP (**B, right**) were measured. (**C**) The relative mRNA levels of SARS-CoV-2 E (**C, left**), N gene (**C, right**), were measured from lung homogenates in indicated groups. The data were expressed as fold changes relative to the non-IAV infection control. (**D**) Histopathologic and immunohistochemical studies were performed with lung slide samples in indicated groups. (**B-D**) Values are mean ± s.d. of three independent results. *P≤0.05, **P≤0.01, ***P≤0.001, ****P≤0.0001.

### IAV components specifically facilitate the entry process of SARS-CoV-2

We further tested if several other viruses on hand had similar effects to promote SARS-CoV-2 infection. To our surprise, neither Sendai virus (SeV) (Fig. 3A), human rhinovirus (HRV3) (Fig. 3B), human parainfluenza virus (HPIV) (Fig. 3C), human respiratory syncytial virus (HRSV) (Fig. 3C) nor human enterovirus 71 (EV71) (Fig. 3C) could stimulate SARS-CoV-2 infection.

**Fig. 3.**
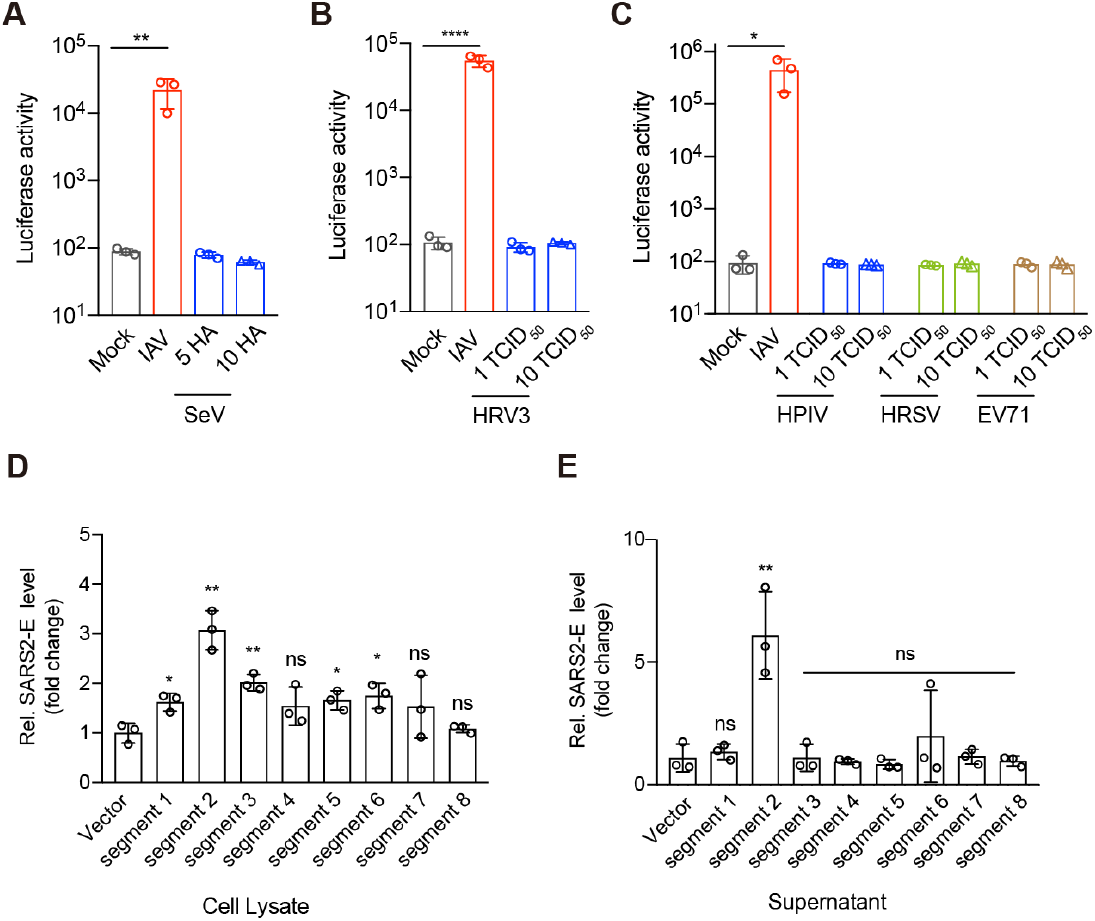
The enhancement of SARS-CoV-2 infection especially responses to IAV. (**A-C**) A549 cells were pre-infected with SeV, HRV3, HPIV, HRSV, or EV71 at indicated doses for 12 hours respectively. Cells were then infected with pSARS-CoV-2 for another 24 hours followed by measuring luciferase activity. (**D and E**) The eight individual segment of WSN were transfected to A549 cells 24 hours ahead of live SARS-CoV-2 infection. Total RNA was extracted from cell lysates (**D**) or supernatant (**E**) to detect the E gene by Taqman-qRT-PCR 48 hours-post-infection. The data were expressed as fold changes relative to the vector control. Values are mean ± s.d. of three independent results. *P≤0.05, **P≤0.01, ***P≤0.001, ****P≤0.0001.

To explore how IAV promotes SARS-CoV-2 infection, we transfected A549 cells with eight individual viral genome segments of IAV to test if any of them could promote SARS-CoV-2 infectivity. The data in Fig. 3D and Fig. 3E showed that IAV segment-2 expression strongly stimulated SARS-CoV-2 multiplication in both SARS-CoV-2-infected cell lysates and supernatant.

### IAV infection induces elevated ACE2 expression

As IAV strongly increased the pseudotyped SARS-CoV-2 infection, we examined the viral entry process. It was reported that the cellular receptor angiotensin-converting enzyme 2 (ACE2)(*13, 14*), together with transmembrane serine protease 2 (TMPRSS2) (*15*), Furin(*16*) and cathepsin L (CatL)(*17, 18*), mediated SARS-CoV-2 viral entry. In IAV-infected cells, we found that the mRNA level of ACE2 and TMPRSS2, but not Furin and CatL were increased around three folds (A549 in Fig. 4A, Calu-3 in Fig. S2). An obvious switch of intracellular ACE2 expression was triggered at 12 h post-IAV-infection (Fig. 4C). In the meantime, influenza NP, Mx1, and ISG54 increased accordingly confirming a successful infection of IAV (Fig. 4B).

**Fig. 4.**
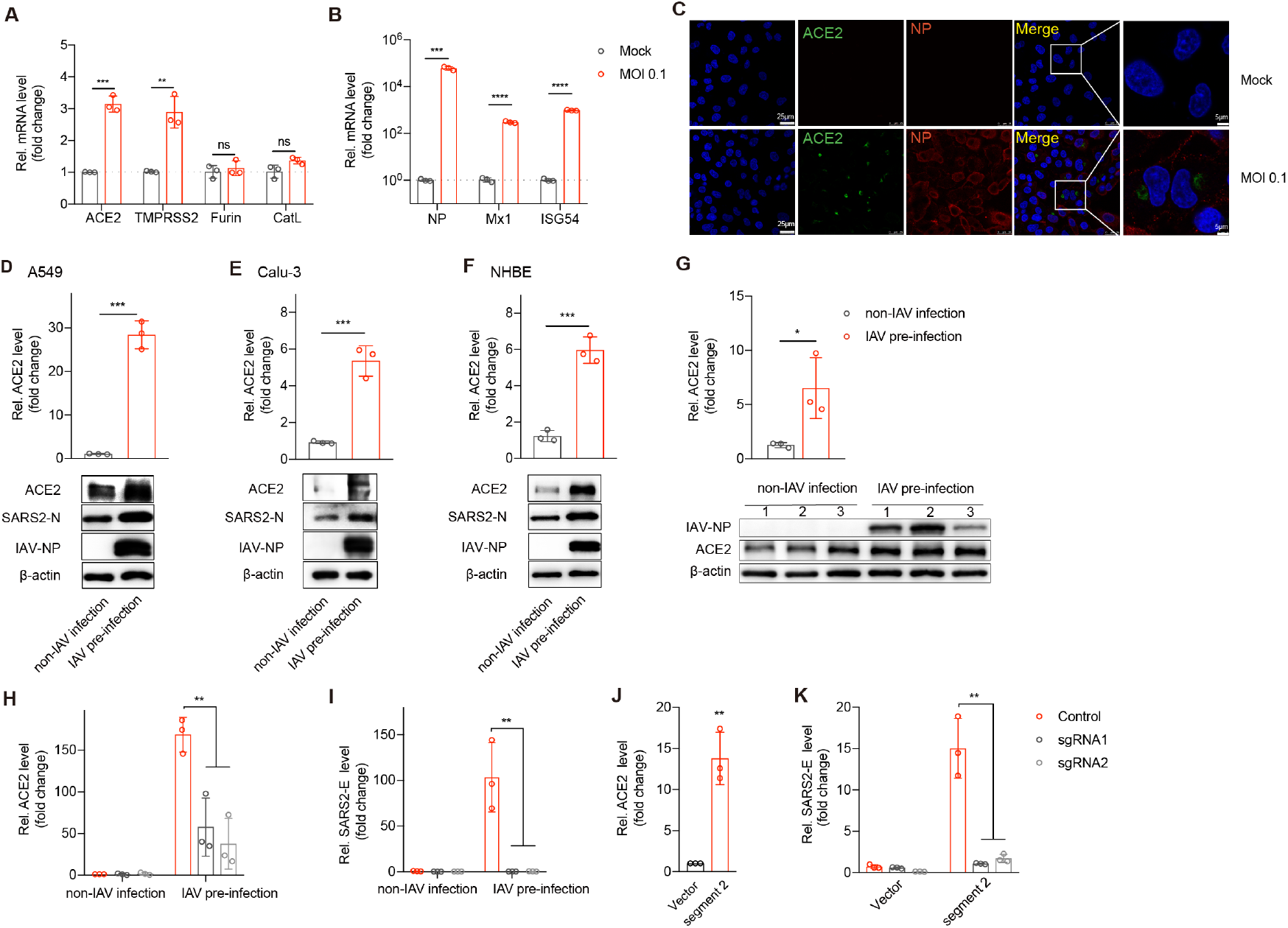
ACE2 is essential for IAV to promote SARS-CoV-2 infection. (**A and B**) A549 cells were mock-infected or infected with WSN at MOI 0.1. At 12 hours post-infection (h.p.i.), total RNAs were extracted from cells, and mRNA of ACE2, TMPRSS2, Furin, CatL (**A**), or mRNA of NP, Mx1, ISG54 (**B**)was evaluated by quantitative real-time PCR (qRT-PCR) using SYBR green method. The data were expressed as fold changes relative to the Mock infections. (**C**) A549 cells were infected with WSN at MOI 0.1. IAV NP proteins (red) and ACE2 (green) were detected by an immunofluorescence assay using a confocal microscope at 12 hours-post-infection. Scale bars were shown. A549 (**D**), Calu-3 (**E**), and NHBE (**F**) cells were pre-infected with WSN at MOI 0.1 for 12 hours. Cells were then infected with live SARS-CoV-2 at MOI 0.01 for another 48 hours. Total RNAs were extracted from cells and mRNA of ACE2 was evaluated by quantitative real-time PCR (qRT-PCR) using SYBR green method. The protein expression levels of ACE2, SARS-CoV-2 N gene, IAV NP, and β-actin were measured by western blot. (**G**) The relative mRNA levels of ACE2 were measured from lung homogenates in indicated groups and the protein expression of IAV NP and ACE2 were detected by western blot accordingly. (**D-G and J**) The data were expressed as fold changes relative to the non-IAV infection control. (**H-K**) To establish ACE2 knock-down cells, A549 cell mixture was transduced with lentivirus encoding CRISPR-Cas9 system with two guide RNAs targeting ACE2 (sgRNA1 and sgRNA2) or control guide RNA respectively. Cells were infected with live SARS-CoV-2 at MOI 0.01 with or without IAV infection under the same procedure as above. The mRNA levels of ACE2 (qRT-PCR) (**H**) and SARS-CoV-2 E gene (Taqman-qRT-PCR) (**I**) expression were detected. (**J**) The mRNA level of ACE2 was detected by qRT-PCR in live SARS-CoV-2-infected cells transfected with either vector of WSN segment-2 respectively. (**K**) The mRNA levels of the SARS-CoV-2 E gene from either vector- or segment2-transfected cells were measured by Taqman-qRT-PCR at 48 hours post-live-SARS-CoV-2-infection in the present of control sgRNA or ACE2 sgRNAs. The data were expressed as fold change relative to non-IAV infection control. Values are mean ± s.d. of three independent results. *P≤0.05, **P≤0.01, ***P≤0.001, ****P≤0.0001.

Interestingly, ACE2 mRNA level increased more dramatically in IAV and SARS-CoV-2 co-infection cells with 28 folds in A549 (Fig. 4D), 5 folds in Calu-3 (Fig. 4E), 6 folds in NHBE (Fig. 4F) respectively. The mRNA and protein levels (Fig. 4G) of ACE2 also increased accordingly in lung homogenates from co-infection mice.

When the cell mixture was transduced by lentivirus coding ACE2-sgRNA to knockdown ACE2 expression (Fig. 4H), the IAV-mediated enhancement of SARS-CoV-2 infection was totally abolished (Fig. 4I). Consist of this, ACE2 mRNA levels increased 13.8-fold in SARS-CoV-2-infected cells expressing segment-2 compared to that in control cells transfected with vector (Fig. 4J). Again, the enhanced SARS-CoV-2 infectivity mediated by segment-2 could be blocked in ACE2 knock-down cells (Fig. 4K).

The data indicated that IAV permitted increased SARS-CoV-2 infection through the up-regulation of ACE2 expression.

### Enhanced SARS-CoV-2 infectivity is independent of IFN signaling

ACE2 was reported to be an interferon-stimulated gene (ISG) in human airway epithelial cells(*19*). IAV infection will also stimulate type I IFN signaling. We, therefore, tested whether the augment of ACE2 expression is dependent on IFN or not. For this, cells were firstly pre-treated with different doses of IFNα (Fig. 5) and IFNγ (Fig. S3 A-C) and then infected with pSARS-CoV-2. The data showed that IFNα could not promote the pSARS-CoV-2 infectivity in A549 cells (Fig. 5A), but rather significantly inhibit pSARS-CoV-2 infectivity in Calu-3 (Fig. 5D) and Huh-7 (Fig. 5G) cells. Compared with the mRNA levels of ISG54 (Fig. 5 B, E, H), the mRNA levels of ACE2 and TMPRSS2 were only mildly increased around 1-3 folds under IFN treatment (Fig. 5 C, F, I). The data indicated that ACE2 could not robustly respond to IFN in these cells, which in turn suggested that ACE2 mediated viral entry was not affected by IFN.

**Fig. 5.**
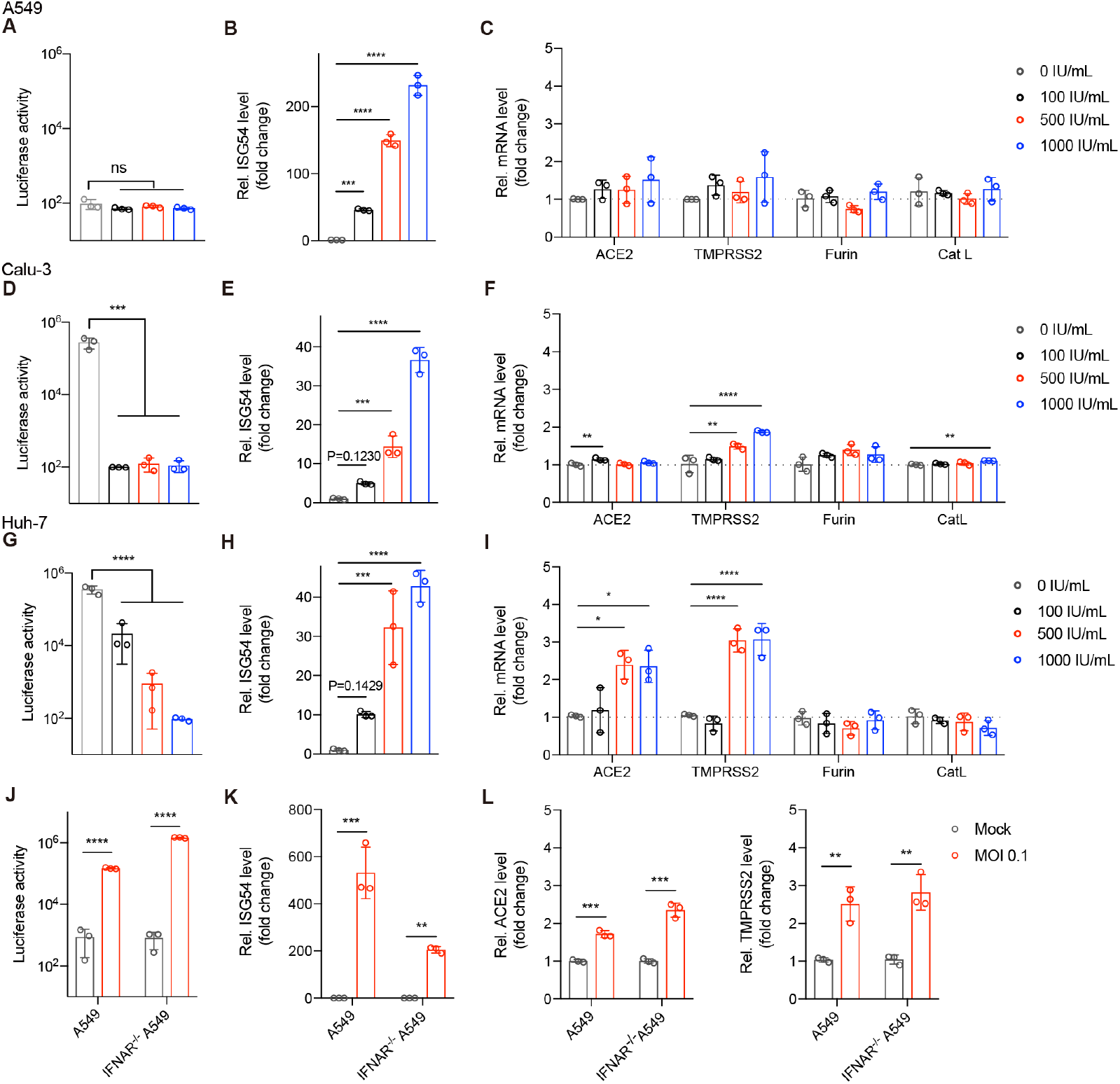
Enhanced SARS-CoV-2 infection is independent of IFN signaling. A549 (**A, B, C**), Calu-3 (**D, E, F**), and Huh-7 (**G, H, I**) cells were pre-treated with indicated doses of IFNα for 12 hours. Cells were then infected with pSARS-CoV-2 for another 24 hours followed by measuring luciferase activity and mRNA expression levels of indicated genes. The data of mRNA levels were expressed as fold changes relative to non-treatment cells. (**J-L**) WT A549, and IFNAR^-/-^A549 cells were infected with W SN at MOI 0.1 for 12 hours, cells were then infected with pSARS-CoV-2 for another 24 hours followed by measuring luciferase activity and mRNA expression levels of indicated genes. P values are from unpaired One-way ANOVA. Values are mean ± s.d. of three independent results. *P≤0.05, **P≤0.01, ***P≤0.001, ****P≤0.0001.

Moreover, in IFNAR^-/-^ A549 cells, the enhanced infectivity of pSARS-CoV-2 under IAV co-infection remained (Fig. 5J). By contrast to the decreased levels of ISG54 in IFNAR^-/-^ A549 cells (Fig. S3D and Fig. 5K), the mRNA levels of ACE2 and TMPRSS2 still increased in IFNAR^-/-^ A549 cells under IAV infection (Fig. 5L). The results strongly suggested that SARS-CoV-2 responded to IAV infection rather than IFN signaling for a favorable viral infection.

## Discussion

Recently, there are many discussions about the possible impacts of the upcoming flu season on the current COVID-19 pandemic. Speculations have been raised that infection of IAV could induce more severe disease for the secondary SARS-CoV-2 infection, or co-infection of these two viruses cause more serious illness. However, no experimental data are available to show the relationship between IAV and SARS-CoV-2 yet. In this study, we provide the first experimental evidence that the pre-infection of IAV strongly promotes SARS-CoV-2 virus entry and infectivity in co-infected cells and animals. It emphasizes that influenza prevention during the SARS-CoV-2 pandemic season is of great importance.

Co-infection of viruses frequently occurs in nature. Some studies showed positive interaction between the dengue virus and the Zika virus via antibody-dependent enhancement(*20*). Other studies showed negative interactions between the common cold virus and SARS-CoV-2 via pre-existing immunity(*21*). By co-infection with IAV and pseudotyped or live SARS-CoV-2, we observed a great enhancement of SARS-CoV-2 infectivity both in cell culture and *in vivo* in infected mice. Such enhancement was associated with the increased expression level of ACE2 which is a major receptor for SARS-CoV-2 to enter a host cell. We detected a 2-3 folds increase in ACE2 mRNA level post-IAV-infection (A549 cells). However, a much higher increase (28 folds) in the ACE2 mRNA level could be detected under IAV and SARS-CoV-2 co-infection. We suspected that IAV infection induced a mild expression of ACE2 to permit SARS-CoV-2 virus entry so that the subsequent multiplication of SARS-CoV-2 would further enhance ACE2 expression in a positive feedback pattern(*19*).

Intriguingly, among all the viruses tested on hand, only IAV but not SeV, HRV3, HPIV, HRSV, or EV71 promoted SARS-CoV-2 infection. The three viruses of HRV3, HPIV, and HRSV are all prevalent pathogens to cause common cold in humans but had no effects on SARS-CoV-2 infectivity. EV71 is a major causative agent for hand-foot-and-mouth disease in young children, but again had little influence on SARS-CoV-2 infection. Furthermore, we confirmed the effects of IAV by H1N1 and H3N2 natural isolates (Fig. S4A), and the infectivity of the current D614G mutant SARS-CoV-2 can also be stimulated by IAV pre-infection (Fig. S4B). The unique feature for IAV to augment SARS-CoV-2 infectivity indicates that the influenza virus is the key pathogen of prevention and control during the current coronavirus pandemic.

Among the eight segments of IAV, segment-2 encoding PB1 promotes ACE2 expression and SARS-CoV-2 infectivity at the highest level. The detailed molecular mechanism underlying PB1 mediated SARS-CoV-2 enhancement needs further study. Nevertheless, the IAV PB1 segment encodes multiple viral proteins including PB1, PB1-F2, PB1-N40 to modulate host cells(*22*). PB1-F2 is a pro-apoptotic factor and can regulate innate immunity(*23*). PB1-N40 interacts with many host factors and contributes to viral pathogenicity(*24*). After all, the fact that the IAV PB1 segment could promote SARS-CoV-2 infection further confirms a unique positive interaction between IAV and SARS-CoV-2.

Importantly, the enhancement phenotype in IAV and SARS-CoV-2 co-infection is independent of IFN signaling. Therefore, influenza vaccination should be recommended to people under the high risk of co-infection. Our findings remind the society that surveillance of co-infection is encouraged in the coming winter. And for sure, social distance and mask-wearing are beneficial to protect people from attacks of either or both the influenza virus.

## Methods

### Cells and viruses

The 293T, A549, Huh-7, MDCK, and Vero E6, WI-38, WI-38 VA-13, and BEAS-2B were obtained from ATCC and maintained in Dulbecco’s modified Eagle’s medium (DMEM; Gibco) supplemented with 10% foetal bovine serum (FBS), Calu-3 (ATCC) was maintained in DMEM supplemented with 20% FBS. NCI-H292(ATCC) was maintained with RPMI-1640 (Gibco) supplemented with 20% FBS. Normal Human Bronchial Epithelial cells (NHBE) cells (ATCC) were maintained in airway epithelial cell basal medium (ATCC PCS300030) supplemented with Bronchial/Tracheal Epithelial Cell Growth Kit (ATCC PCS-300-040). All cells were incubated at 37 °C, 5% CO2.

The A/WSN/33 virus was generated by reverse genetics as previously described(*25*). H1N1(A/Sichuan/01/2009) and H3N2 (A/Donghu/312/2006) were kindly provided by the Influenza Center in China CDC. Human rhinovirus (HRV3), human respiratory syncytial virus (HRSV), or human enterovirus 71 (EV71) were purchased from ATCC and stocked accordingly. The human parainfluenza virus (HPIV) was obtained from Prof. MingZhou Chen, Wuhan University. Sendai virus (SeV) was provided by Prof. Tianxian Li, Wuhan Institute of Virology. The SARS-CoV-2 live virus (strain IVCAS 6.7512) was provided by the National Virus Resource, Wuhan Institute of Virology, Chinese Academy of Sciences.

### Plasmids and transfection

The SARS-CoV-2-S-Δ18 expressing plasmid was a gift from Prof. Ningshao Xia, Xiamen University. The eight WSN viral segments in pHW2000 plasmid were kindly provided by Prof. Hans Klenk, Marburg University. The DNA transfection reagent Fugene HD was purchased from Promega and the transfection was performed according to manuscript procedures.

### Pseudotype virus production

The pseudotyped VSV-ΔG viruses expressing either luciferase reporter or mCherry reporter were provided by Prof. Ningshao Xia, Xiamen University. To produce pseudotyped VSV-ΔG-Luc/mCherry bearing SARS-CoV-2 spike protein (pSARS-CoV-2), Vero E6 cells were seeded in 10 cm dish and transfected simultaneously with 15 μg SARS-CoV-2-S-Δ18 plasmid by Lipofectamine 3000 (Thermo). Forty-eight hours post-transfection, 150 μl pseudotyped VSV-ΔG bearing VSV-G protein were used to infect Vero E6 cells. Cell supernatants were collected after another 24 hours clearing from cell debris by centrifugation at 3000rpm for 6 minutes, aliquoted and stored at - 80 °C.

### Luciferase-based cell entry assay

Target cells were seeded in 48-well plates and inoculated, in triplicate, with equivalent volumes of pseudotyped virus stocks with 1:5 dilution in DMEM (3% FBS) with or without IAV pre-infection. At 24 h post-pseudotype-infection, the luciferase activities were measured with the Luciferase Assay System (Promega E4550).

### Virus infection and IFN treatment

For IAV infection, cells were washed with PBS and then incubated with viruses at different MOIs (from 0.01 to 1) in infection medium (DMEM, supplemented with 2% FBS, 1% penicillin/streptomycin) at 37 °C, 5% CO2.

For SARS-CoV-2 infections, cells were incubated with SARS-CoV-2 live virus at MOI of 0.01 in infection medium (DMEM, 1% penicillin/streptomycin) and incubated at 37 °C, 5% CO2 for 1 hour with or without 12 h IAV pre-infection (MOI 0.1). Cells were then washed with PBS two times and then incubated in culture medium (DMEM, supplemented with 5% FBS, 1% penicillin/streptomycin) at 37 °C, 5% CO2 for 48 hours.

For SeV, HRV3, HPIV, HRSV, or EV71 infection, cells were washed with PBS and then incubated with indicated viruses in infection medium (DMEM, supplemented with 3% FBS, 1% penicillin/streptomycin) and incubated at 37 °C, 5% CO2 for 12 hours.

For IFN treatment, recombinant human IFNα 2a (Beyotime, P5646) and IFNγ (Beyotime, P5664) were dissolved in 0.1% BSA and diluted in DMEM with 10% FBS, and then admitted to cells for 12 hours at indicated doses.

### Real-time reverse-transcriptase–polymerase chain reaction

The mRNA levels of indicated genes were quantified by quantitative PCR with reverse transcription (qRT–PCR). Purified RNAs extracted by TRIzol (Invitrogen™,15596018) were subjected to reverse transcription with oligo dT primer (using Takara cat#RR037A Kit), and then the corresponding cDNAs were quantified using Hieff qPCR SYBR Green Master Mix (Yeason). Thermal cycling was performed in a 384-well reaction plate (ThermoFisher, 4343814). Gene-specific primers were shown in Supplementary Table 1. All the mRNA levels were normalized by β-actin in the same cell.

The relative number of SARS-CoV-2 viral genome copy number were determined using Taqman RT-PCR Kit (Yeason). To acutely quantify the absolute number of SARS-CoV-2 genome, a standard curve by measuring the SARS-CoV-2 N gene constructed in the pCMV-N plasmid was applied. All the SARS-CoV-2 genome copy numbers were normalized by GADPH in the same cell.

### Western blot analysis

For western blots, cells were lysed in RIPA buffer on ice for 30 minutes and were separated by sodium dodecyl sulfate-polyacrylamide gel electrophoresis (SDS–PAGE) and subjected to western blot analysis. For mice experiments, half lung tissue from each mouse was homogenized in PBS followed by boiling in SDS lysis buffer (GE) at 100°C, 30 minutes. Rabbit monoclonal antibody against ACE2 (Abclonal, A4612, 1:1000), mouse monoclonal antibody against SARS-CoV Nucleoprotein (Sino Biological, 40143-MM05, 1:1000), anti-actin (Abclonal, 1:1000), were purchased commercially. The anti-influenza virus-NP was kindly provided by Prof. Ningshao. Xia. Peroxidase-conjugated secondary antibodies (Antgene, 1: 5000) were applied accordingly followed by image development with Chemiluminescent HRP Substrate Kit (Millipore Corporation).

### Immunofluorescence

A549 cells were fixed and incubated with primary antibodies. The primary antibodies used in this study were rabbit polyclonal antibody against ACE2 for immunofluorescence (Sino Biological, 10108-T26) and anti-influenza virus-NP (kindly provided by Prof. Ningshao Xia). The Alexa Fluor dye-conjugated secondary antibodies (Alexa Fluor R488, Invitrogen; Alexa Fluor M555, Invitrogen) and DAPI (Beyotime, C1002), were admitted afterward according to standard protocols. Cell imaging was performed on a Leica TCS SP8 confocal laser scanning microscope (Leica).

### ACE2 knocking-down cells

Two sgRNAs targeting the hACE2 gene were designed under the protocol in http://chopchop.cbu.uib.no and commercially synthesized to clone in lenti-Cas9-blast vector (kindly provided by Prof. Hongbing Shu). The control sgRNA lentivirus construct was also provided by Prof. Hongbing Shu. In brief, A549 cells were plated at 6-well plates and transduced with lentivirus encoding CRISPR-Cas9 system including either ACE2 sgRNA or control sgRNA. The cell mixtures were selected by blasticidin for one week to obtain ACE2 knocking-down cells. The gene knocking-down efficiencies were confirmed by measuring the ACE2 mRNA level through qRT-PCR analysis.

### Mice

The K18 hACE2 transgenic mice purchased from Gempharmatech were housed in ABSL-3 pathogen-free facilities under 12-h light-dark cycles with access to food and water. Mice were male, age-matched, and grouped for SARS-CoV-2 infection or IAV and SARS-CoV-2 co-infection. At day 0, mice were intranasally infected with PBS or 2000 PFU of WSN respectively, and then both groups were intranasally infected with 3 x10^5^ PFU of SARS-CoV-2 at Day 2. Another two days later, mice were sacrificed to determine viral loads and submitted to histological assay.

### Histology

Lung tissue from infected mice was dissected at Day 2 post-SARS-CoV-2-infection, fixed, and stained using a standard H&E procedure. The slides were scanned and analyzed by the Wuhan Sci-Meds company. The representative images from three mice in each group were shown.

### Statistical analysis

If not indicated otherwise, Student’s t-test was used for two-group comparisons. The *p-value < 0.05, **p-value < 0.01, ***p-value < 0.001 and ****p-value < 0.0001 were considered significant. Unless otherwise noted, error bars indicated as mean values with standard deviation of at least three biological experiments.

## Author contributions

K.X. and K.L. conceived the project and designed the experiments. L.B., J. D., M.G., X.W., Z. H., Z. Z., and YC. Z. coordinated the live SARS-CoV-2 study and performed animal infection experiments. YL. Z. and S. L. conducted pseudotyped virus infection experiments, IFN treatment experiments, and data analysis. L.B., J. D. solved the Immunofluorescence, Histopathologic and Immunohistochemical studies. X. L. performed SeV, HRV3, HPIV, HRSV, EV71 infection experiments. YL. Z and X. L. generated the mutant virus and performed the related test. L.B., S. L, J. D., and X. L. repeated the key experiments in infected cells. X. S., Q.L., D. N., M.X., K.S., J.Y., W.Z., Z. T., M. T., Y. Z., C.S., M. D., L.Z., Y.C., and H.Y provided technical supports and the materials. L. D. carried out ACE2 knock-out cells and related analysis. K.X., K.L., S. L, and YL. Z wrote the manuscript with inputs from all the remaining authors. We also thank our group members of the SARS-CoV-2 working group in the State Key Laboratory of Virology, Wuhan University, who work tightly together during this new virus pandemic for their research spirits and courage. We are grateful to Taikang Insurance Group Co., Ltd, Beijing Taikang Yicai Foundation, and Special Fund for COVID-19 Research of Wuhan University for their great supports of this work.

---

**Fig. S1.**
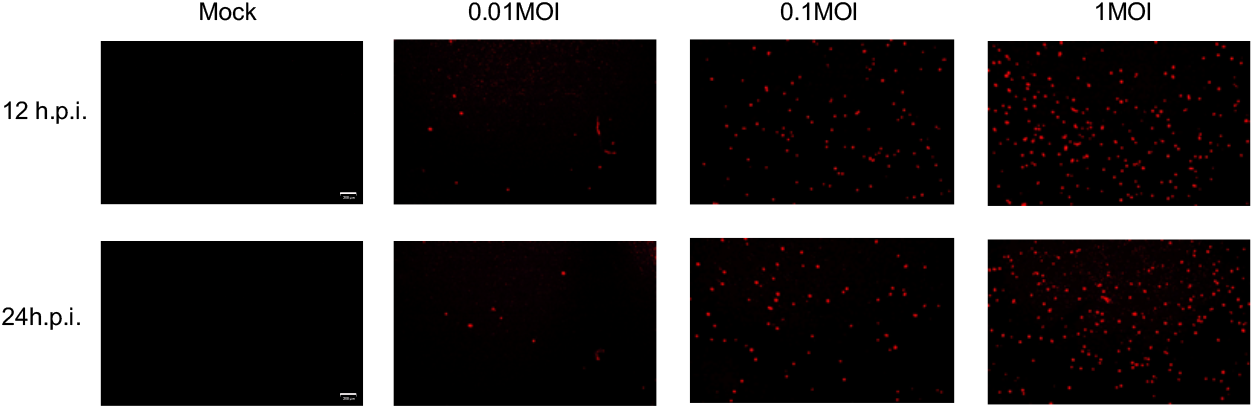
IAV facilitates the entry process of pSARS-CoV-2 (relate to Fig.1). A549 cells were infected with A/WSN/33 at indicated MOIs. At 12, 24 hours post-IAV-infection, cells were infected with pSARS-CoV-2 with mCherry reporter for another 24 hours. Scale bars, 200 μm.

**Fig. S2.**
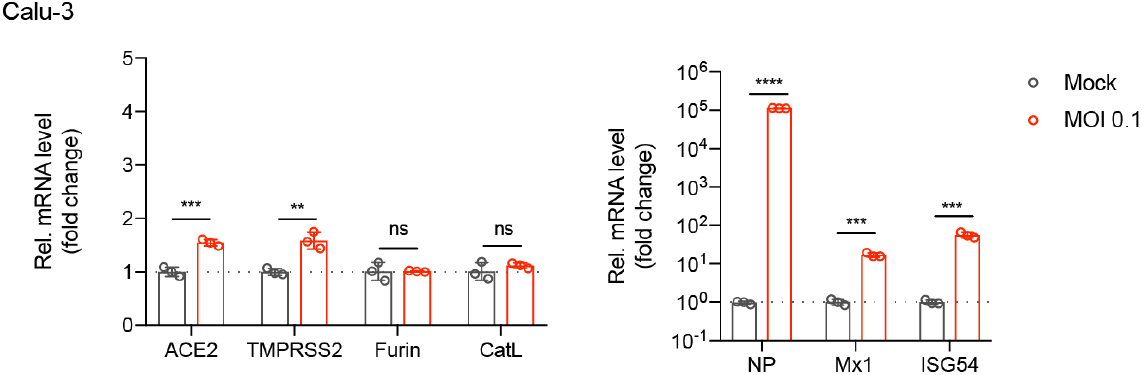
IAV infection induces elevated ACE2 expression (relate to Fig.4). Calu-3 cells were mock-infected or infected with WSN at MOI of 0.1. At 12 hours h.p.i., total RNAs were extracted from cells, and mRNA of ACE2, TMPRSS2, Furin, CatL, NP, Mx1, and ISG54 were evaluated by qRT-PCR using the SYBR green method. The data were expressed as fold changes relative to the Mock infections. Values are mean ± s.d. of three independent results. *P≤0.05, **P≤0.01, ***P≤0.001, ****P≤0.0001.

**Fig. S3.**
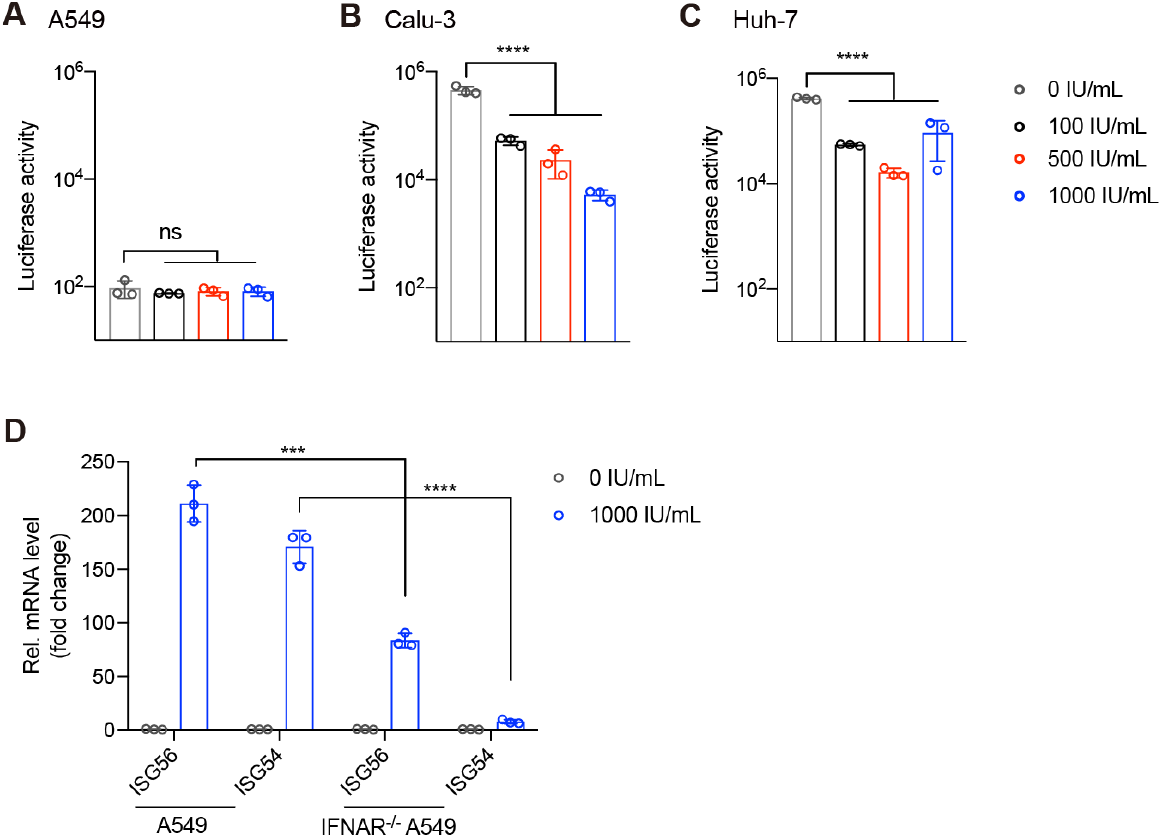
Enhanced SARS-CoV-2 infection is independent of IFN signaling (relate to Fig.5). A549 (**A**), Calu-3 (**B**), and Huh-7 (**C**) cells were pre-treated with indicated doses of IFNγ for 12 hours. Cells were then infected with pSARS-CoV-2 for another 24 hours followed by measuring luciferase activity. (**D**) WT, and IFNAR^-/-^A549 cells were treated with IFNα at 1000 IU/mL for 12 hours, and the mRNA expression levels of indicated genes were measured. Values are mean ± s.d. of three independent results. (**A-C**) P values are from unpaired One-way ANOVA. *P≤0.05, **P≤0.01, ***P≤0.001, ****P≤0.0001.

**Fig. S4.**
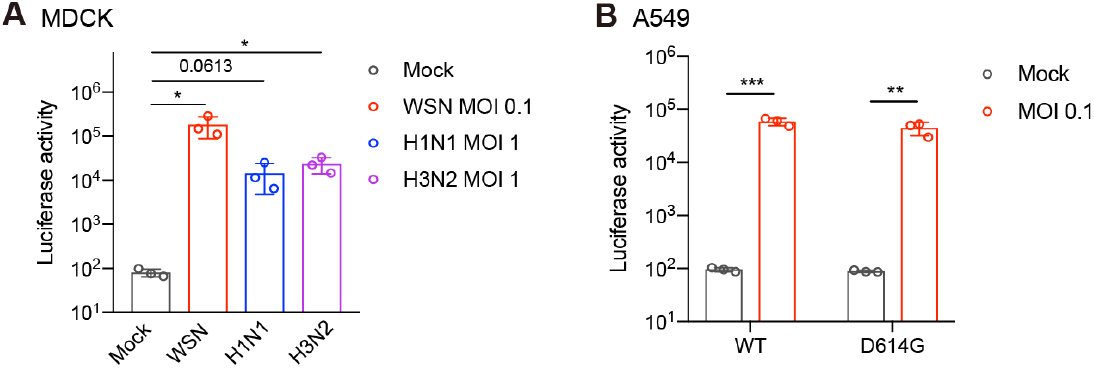
IAV facilitates viral entry of WT or mutant SARS-CoV-2. (**A**) MDCK cells were pre-infected with WSN (MOI=0.1), H1N1(MOI=1), or H3N2 (MOI=1) for 12 hours and were then infected with pSARS-CoV-2 for another 24 hours followed by measuring luciferase activity. (**B**) A549 cells were pre-infected with WSN at MOI 0.1 for 12 hours and were then infected with D614G mutant pSARS-CoV-2 for another 24 hours followed by measuring luciferase activity. Values are mean ± s.d. of three independent results. *P≤0.05, **P≤0.01, ***P≤0.001.

**Supplementary Table 1.**
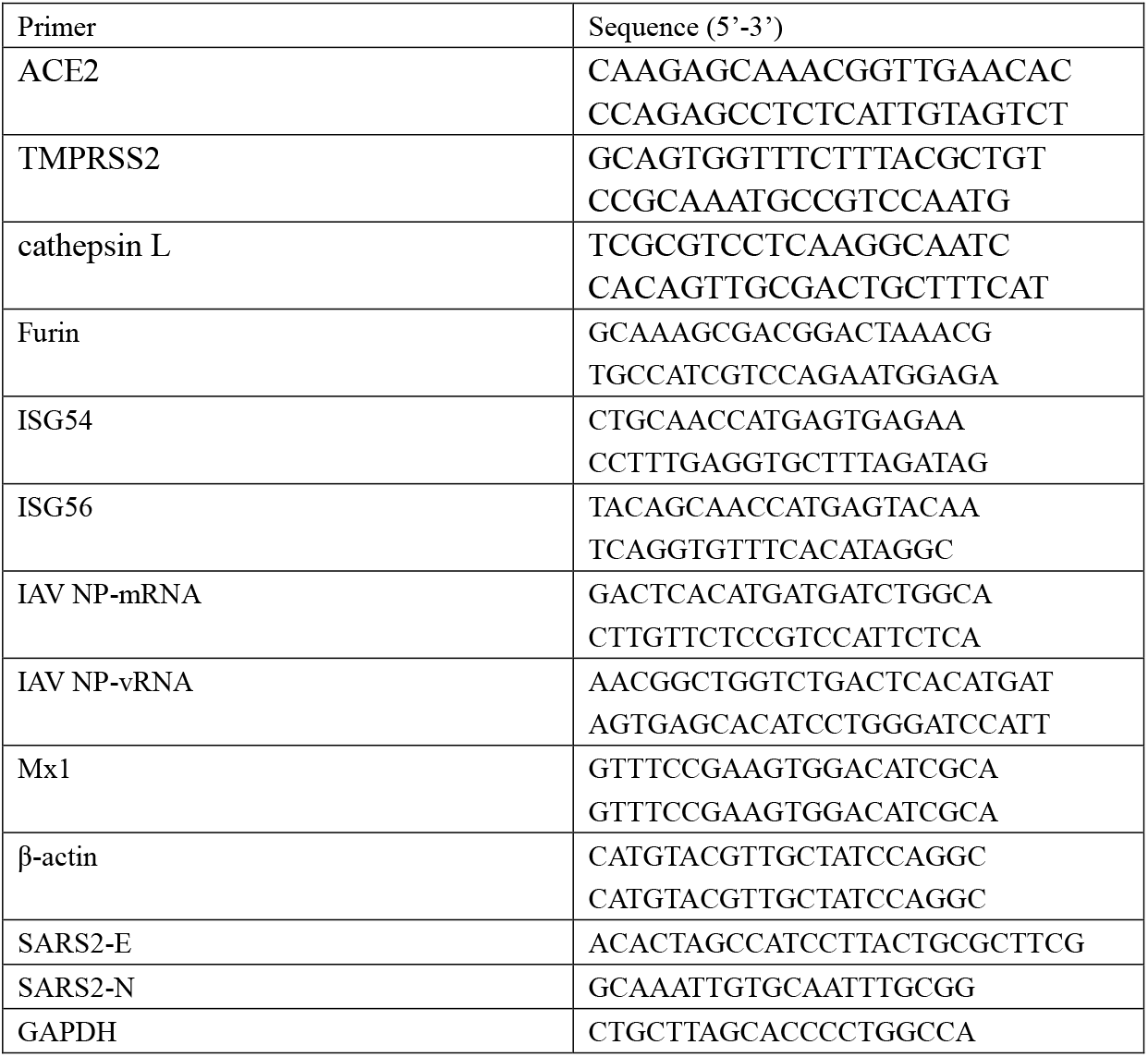
Primers used in this paper.

